# Functional and Structural Characterization of OXA-935, a Novel OXA-10-family β-lactamase from *Pseudomonas aeruginosa*

**DOI:** 10.1101/2021.09.20.461173

**Authors:** Nathan Pincus, Monica Rosas-Lemus, Samuel Gatesy, Ludmilla Shuvalova, Joseph Brunzelle, George Minasov, Karla Satchell, Marine Lebrun-Corbin, Egon Ozer, Alan Hauser, Kelly Bachta

## Abstract

Resistance to antipseudomonal penicillins and cephalosporins is often driven by the overproduction of the intrinsic β-lactamase AmpC. However, OXA-10-family β-lactamases are a rich source of resistance in *Pseudomonas aeruginosa*. OXA β-lactamases have a propensity for mutation leading to extended spectrum cephalosporinase and carbapenemase activity. In this study, we identified isolates from a subclade of the multidrug-resistant (MDR) high risk clonal complex CC446 with resistance to ceftazidime. Genomic analysis revealed that these isolates harbored a plasmid containing a novel allele of *bla*_OXA-10_, named *bla*_OXA-935_, which was predicted to produce an OXA-10 variant with two amino acid substitutions: an aspartic acid instead of glycine at position 157 and a serine instead of phenylalanine at position 153. The G157D mutation, present in OXA-14, is associated with resistance to ceftazidime. Deletion of *bla*_OXA-935_ restored sensitivity to ceftazidime and susceptibility profiling of *P. aeruginosa* laboratory strains expressing *bla*_OXA-935_ revealed that OXA-935 conferred ceftazidime resistance. To better understand the impact of the variant amino acids, we determined the crystal structures of OXA-14 and OXA-935. In OXA-14, one of two monomers contained the canonical carbamylated lysine-70 (K70). In contrast, both monomers of OXA-935 were decarbamylated at K70, and the F153S mutation conferred increased flexibility to the omega (Ω) loop. Compared to OXA-14, the catalytic efficiency of OXA-935 for nitrocefin was significantly reduced. Amino acid changes that confer extended spectrum cephalosporinase activity to OXA-10-family β-lactamases are concerning given rising reliance on novel β-lactam/β-lactamase inhibitor combinations such as ceftolozane-tazobactam and ceftazidime-avibactam to treat MDR *P. aeruginosa* infections.

## INTRODUCTION

Infections caused by multidrug-resistant (MDR) and extensively drug-resistant (XDR) organisms are an increasing threat to public health. Leading the way are infections by drug-resistant gram-negative bacteria such as *Pseudomonas aeruginosa, Acinetobacter baumannii* and members of the *Enterobacteriaceae* family. Of particular interest is the human pathogen, *P. aeruginosa*, which is responsible for diverse infections including bacteremia, pneumonia, urinary tract and skin and soft tissue infections. At baseline, *P. aeruginosa* harbors a large armamentarium of antimicrobial resistance mechanisms including a chromosomally encoded cephalosporinase, AmpC, and basal and inducible antibiotic efflux pumps. *P. aeruginosa* is notable for the frequent acquisition of exogenous genetic material in the form of plasmids that carry large mobile genetic elements (e.g. integrons, transposons, ICE elements) that can contain series of antimicrobial resistance genes (1, 2). One such element, the class 1 integron in1697, was recently identified as part of a novel antimicrobial resistance (AMR) plasmid discovered in ST298*, a subclade within the globally-distributed *P. aeruginosa* high-risk clonal complex 446 (CC446) (1). Characterization of in1697 revealed the presence of several resistance gene cassettes including genes for sulfonamide, quaternary ammonium compounds and aminoglycoside resistance. In most strains, in1697 also contained the β-lactamase gene, *bla*_*OXA-10*_.

OXA-10 (PSE-2) is a class D β-lactamase originally described in the late 1970s (3, 4) that confers resistance to cefotaxime and ceftriaxone by not ceftazidime (5). Natural and laboratory-selected variants of OXA-10 have been identified, many of which differ in the spectrum of β-lactams that they hydrolyze (6). One such variant, OXA-14, contains a single amino acid change (glycine to aspartate at position 157, G157D) from the OXA-10 parent. The OXA-14 variant has been implicated in clinically significant resistance to ceftazidime (6, 7). In *P. aeruginosa*, variations in OXA-10-type enzymes are a rich source of expanding AMR, as evidenced by several recently identified members of the class, OXA-40, OXA-198, OXA-655 and OXA-656, that confer resistance to carbapenems (8-11).

Although the isolates within this collection of ST298* *P. aeruginosa* strains were typically susceptible to ceftazidime, we noted three isolates that were resistant to this antibiotic (1). Interestingly, these three isolates also contained AMR integrons in which the *bla*_*OXA-10*_ gene contained two mutations that were predicted to result in a glycine to aspartic acid substitution at position 157 (similar to OXA-14) and a phenylalanine to serine substitution at position 153. In the present study, we characterized this novel OXA-10 variant, which we designate “OXA-935”. We determined the crystal structures of both OXA-935 and OXA-14. The structures of OXA-935 and OXA-14 shared most of the structural features previously described for OXA-10 (12); however, OXA-935 contained a second amino substitution, phenylalanine to serine at position 153 (F153S), that increased the flexibility of the Ω-loop resulting in the loss of critical carbamylation at the active site residue lysine 70 (K70).

## RESULTS

### Identification of OXA-935, a novel class D, OXA-10 family β-lactamase

In a recent study, we described a prolonged epidemic of ST298* XDR *P. aeruginosa* at a single academic center (1). Many of these strains possessed sequence aligning to a novel AMR plasmid (pPABL048) harboring the integron, in1697. This plasmid, originally described in the strain PABL048, confers resistance to anti-pseudomonal penicillins likely due to the presence of *bla*_OXA-10_ in the integron. We noted that three isolates (PS1793, PS1796, and PS1797) showed high levels of ceftazidime resistance not present elsewhere in the collection. Additionally, they also showed non-susceptibility to ceftazidime-avibactam and ceftolozane-tazobactam by disc diffusion. These three isolates were closely related and found to possess single nucleotide variants (SNVs) in the plasmid-borne *bla*_OXA-10_ gene leading to G157D and F153S amino acid substitutions in OXA-10. These isolates also harbored deletions of amino acids 2-30 in AmpD (1). AmpD negatively regulates the chromosomally encoded cephalosporinase, AmpC and mutations in AmpD are linked to overexpression of AmpC and increased resistance to cephalosporins (13, 14). In this study, we explored the molecular mechanism of ceftazidime resistance in PS1793, PS1796 and PS1797 with a focus on their OXA-10-variant β-lactamase.

We performed long-read sequencing of PS1793 and used this, along with previously-generated Illumina short reads (1), to assemble a complete genome sequence for PS1793. This yielded a 7.4 Mb genome, consisting of a 6,868,713 bp circular chromosome and 3 circular plasmids. Surprisingly, the AMR plasmid pPABL048 described in our previous study aligned to two separate plasmid sequences in PS1793 (the 318,215 bp PS1793_p1 and 113,189 bp PS1793_p2) with approximately 19 kb of overlapping sequence present on both plasmids (Fig. 1A-D). No additional sequence was present in these plasmids, suggesting that pPABL048 may, in fact, be a hybrid plasmid that contains more than one set of replication and partitioning machinery. This would be similar to other plasmids within this family such as pOZ176 that possesses both the IncP-2 system and the uncharacterized replication gene described in our previous study (1, 15). The AMR integron is present in the larger PS1793_p1 plasmid. PS1793 also harbors an additional 69,506 bp plasmid (PS1793_p3) that was not previously described in PABL048 (Figure 1A-D). However, following a BLAST search, portions of this third plasmid aligned with the PABL048 chromosome (a 10 kb fragment aligned with 99% identity and a 2.3 kb fragment aligned with 76% identity) and shared homology with other *Pseudomonas* genus plasmids. Based on alignments with the PS1793 complete genome, PS1796 was identical (with 0 chromosomal or plasmid SNVs) and PS1797 differed by a single chromosomal SNV (Table S1).

**FIGURE 1.**
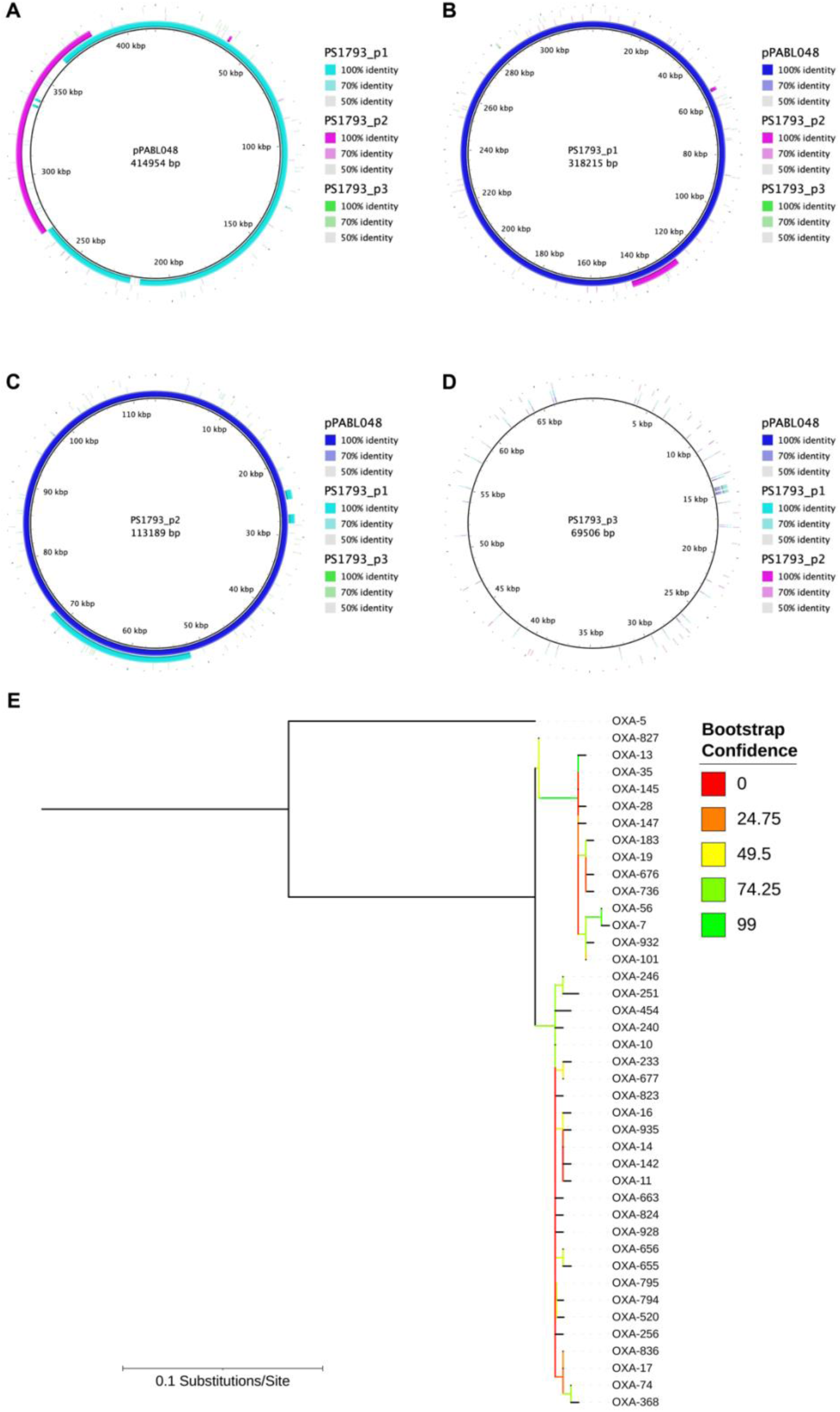
Comparison of the plasmids found in *P. aeruginosa* PS1793 with pPABL048 and phylogenetic relationships between OXA-10-like family serine β-lactamases. **(A)** Alignment of pPABL048 (blue) with the three plasmids present in PS1793, p1 (teal), p2 (magenta) and p3 (green). **(B, C)** Alignment revealed that two separate plasmids, p1 and p2 shared substantial sequence overlap with pPABL048. PS1793_p1 and p2 also share an overlapping 19 kb region. **(D)** The 69 kb PS1793_p3 plasmid did not share substantial sequence with the previously characterized pPABL048 plasmid. **(E)** Maximum likelihood phylogenetic tree of OXA-10-family serine β-lactamases. Tree is rooted at OXA-5, which was included as an outgroup. Bootstrap confidence is indicated by color. Analysis revealed two major families with OXA-10 and OXA-7 as their earliest-identified members.

Comparison of the *bla*_OXA-10_ variant seen in PS1793, PS1796 and PS1797 to the NCBI database revealed that this allele had not been previously described. As such, it was assigned the name *bla*_OXA-935_ (RefSeq ID WP_141989064.1). When its predicted protein sequence was compared to other protein sequences within the OXA-10 family, the G157D substitution was present in multiple homologues, including OXA-11 and OXA-14, which are known to confer extended spectrum resistance to ceftazidime (7, 16). The F153S substitution was unique to OXA-935, although OXA-795 possesses a deletion at positions 153 and 154 (Fig. S1). OXA-795 also confers ceftazidime resistance without an accompanying G157D substitution (17). Phylogenetic analysis of these OXA-10 family proteins showed that they are divided into two major groups classified by their first identified and earliest-named members, OXA-7 and OXA-10. OXA-935 belongs to the OXA-10 subgroup and is most closely related to OXA-11, OXA-14, OXA-16, and OXA-142, although most branches within each group have low bootstrap confidence, likely secondary to limited sequence variability (Fig. 1E).

We next sought to determine the prevalence of *bla*_OXA-10_ family genes within the *Pseudomonas* genus. By screening 9799 genomes, we found that *bla*_OXA-10_-like genes were most common in *P. aeruginosa* but were also present in other species including *P. stuzeri* and *P. putida* (Table S2). The *bla*_OXA-10_ allele was present in a diverse set of species and *P. aeruginosa* STs while other variants were more likely to be limited to a few or a single ST. Thus far, *bla*_OXA-935_ has only been detected in the three ST298* *P. aeruginosa* isolates described in this study (Table S3).

### Expression of OXA-935 confers ceftazidime resistance

After identification of PS1793, PS1796, and PS1797 as the only three isolates in our cohort of ST298* *P. aeruginosa* with resistance to ceftazidime, we sought to characterize the role of the novel beta-lactamase allele, *bla*_OXA-935,_ in ceftazidime resistance. Given that OXA-935 shared the G157D variation with OXA-14 and that expression of OXA-14 has been previously linked to ceftazidime resistance (6, 7), we hypothesized that the novel beta-lactamase allele, *bla*_*OXA*-935,_ conferred ceftazidime resistance on our clinical isolates despite the co-existing truncation of AmpD. Resistance of PS1793, PS1796, and PS1797 was confirmed using the microbroth dilution (MBD) method (Table 1). All three isolates demonstrated similarly high levels of resistance to ceftazidime with MICs of 64 µg/mL. Deletion of *bla*_OXA-935_ from the AMR plasmid of PS1793, PS1796 and PS1797 resulted in a reduction in the ceftazidime MIC to 8 µg/mL, 2 µg/mL and 2 µg/mL, respectively, yielding susceptible phenotypes for all three strains. Deletion of *bla*_OXA-935_ also made the strains susceptible to the 4^th^ generation cephalosporin, cefepime. Previously, when we deleted the entire pPABL048 plasmid from related strains in CC446, we detected the loss of resistance to aztreonam and piperacillin/tazobactam (1). Interestingly, deletion of *bla*_OXA-935_ resulted in minor reductions in the MICs to aztreonam and piperacillin/tazobactam, but did not restore susceptibility. This suggests that there likely are compensatory mechanisms that govern resistance to aztreonam and piperacillin/tazobactam. We did not observe any differences in susceptibility to meropenem, suggesting that OXA-935 is unlikely to contribute to carbapenem resistance. Other than alterations in MICs to cephalosporins, all three Δ*bla*_OXA-935_ deletion strains maintained resistance to gentamicin and ciprofloxacin and susceptibility to colistin. Taken together, these results confirm that ceftazidime resistance in PS1793, PS1796 and PS1797 is driven primarily by the presence of *bla*_OXA-935._

**TABLE 1.** Minimal Inhibitory Concentrations (MICs)

| Strain | MICs (µg/mL) |  |  |  |  |  |  |  |
| --- | --- | --- | --- | --- | --- | --- | --- | --- |
|  | Gent | Cipro | Col | Mero | Pip/Tazo | Az | Cep | Ctz |
| PS1793 | >128 – ns | 32 – ns | 1 | 16 – ns | 64 – ns | 16 - ns | 16 - ns | 64 - ns |
| PS1793 $\Delta bla_{oxa935}$ | >128 – ns | 32 – ns | 0.5 | 16 – ns | 32 – ns | 16 - ns | 8 | 8 |
| PS1796 | >128 – ns | 32 – ns | 1 | 16 – ns | 64 – ns | 16 - ns | 16 - ns | 64 - ns |
| PS1796 $\Delta bla_{oxa935}$ | >128 – ns | 16 – ns | 0.25 | 8 – ns | 32 – ns | 16 – ns | 4 | 2 |
| PS1797 | >128 – ns | 32 – ns | 1 | 16 – ns | 128 - ns | 16 – ns | 32 - ns | 64 – ns |
| PS1797 $\Delta bla_{oxa935}$ | >128 – ns | 16 – ns | 0.25 | 8 – ns | 32 – ns | 16 – ns | 4 | 2 |

Gent: gentamicin, Cipro: ciprofloxacin, Col: Colistin, Mero: Meropenem, Pip/Tazo: Piperacillin-Tazobactam, Az: Aztreonam, Cep: Cefepime, Ctz: Ceftazidime ns: non-susceptible (intermediate and resistant); Clinical Laboratory Standards Institute, MIC Interpretive Standards (µg/mL), 2018.

To evaluate the impact of the G157D variant in OXA-14 and the G157D, F153S variants in OXA-935 on β-lactam resistance, *bla*_OXA-10,_ *bla*_OXA-14,_ *and bla*_OXA-935_ were individually cloned into a plasmid and expressed in *P. aeruginosa* PAO1 and PA14 strain backgrounds. Comparative MIC data for PAO1 and PA14 expressing OXA-10, OXA-14, and OXA-935 are shown in Table 2. Compared to strains expressing a vector control, the expression of OXA-14 resulted in a 16-fold increase in the ceftazidime MIC in PAO1 and an 8-fold increase in PA14, consistent with previous reports (6, 7, 17). Expression of OXA-935 resulted in a 16-fold increase in ceftazidime MIC in PAO1 and a 32-fold increase in PA14, providing evidence that both OXA-14 and OXA-935 confer resistance to ceftazidime, but that expression of OXA-935 has a greater impact on MIC. The detailed molecular mechanism of the differential impact of OXA-14 and OXA-935 expression on ceftazidime resistance in various PA strain backgrounds requires more study. Compared to strains containing an empty vector, expression of OXA-10 resulted in an 8-fold increase in the piperacillin-tazobactam MIC in PAO1 and a 16-fold increase in PA14. Expression of OXA-14 resulted in a 4-fold increase the MIC to piperacillin-tazobactam in both strains while expression of OXA-935 resulted in no increase in the MIC to piperacillin-tazobactam. The impact on cefepime susceptibility was more uniform in that all three proteins, OXA-10, OXA-14, and OXA-935, resulted in a 4-fold increase in MIC when expressed in both strain backgrounds. These results confirm previous findings indicating that OXA-10 has activity against piperacillin-tazobactam but not ceftazidime. In contrast, OXA-935 is not appreciably active against piperacillin-tazobactam in these strain backgrounds but has substantial activity against ceftazidime. OXA-14 has an intermediate resistance phenotype.

**TABLE 2.** MICs for PAO1 and PA14 producing OXA-10, OXA-14, and OXA-935

| Strain | MICs (µg/mL) |  |  |  |
| --- | --- | --- | --- | --- |
|  | Pip/Tazo | Az | Cep | Ctz |
| PAO1 + pPSV37 | 2 | 4 | 1 | 1 |
| PAO1 + pPSV37-*oxa10* | 16 | 8 | 4 | 1 |
| PAO1 + pPSV37-*oxa14* | 8 | 8 | 4 | 16 – ns |
| PAO1 + pPSV37-*oxa935* | 2 | 4 | 4 | 16 – ns |
| PA14 + pPSV37 | 2 | 4 | 1 | 2 |
| PA14 + pPSV37-*oxa10* | 32 – ns | 8 | 4 | 4 |
| PA14 + pPSV37-*oxa14* | 8 | 8 | 4 | 16 – ns |
| PA14 + pPSV37-*oxa935* | 2 | 8 | 4 | 64 – ns |

Pip/Tazo: Piperacillin-Tazobactam, Az: Aztreonam, Cep: Cefepime, Ctz: Ceftazidime ns: non-susceptible (intermediate and resistant); Clinical Laboratory Standards Institute, MIC Interpretive Standards (µg/mL), 2018.

### The amino acid change F153S introduced high flexibility into the Ω-loop of OXA-935

OXA-935 differs from OXA-14 in a single amino acid substitution in the protein’s Ω-loop at position 153 (Fig. 1A). Changes in this loop in multiple classes of β-lactamases are known to increase its flexibility and therefore increase substrate access to the active site (12, 18, 19), which could explain increased activity against bulky substrates such as ceftazidime. Since structures of OXA-14 and OXA-935 were unavailable to corroborate this hypothesis, we determined the crystal structure of OXA-14 and OXA-935. We were able to determine apo structures for both enzymes (Fig. 2, Table S4), but not in complex with ceftazidime. Both protein structures contained two chains in the asymmetric unit, corresponding to a dimer as previously described for OXA-10 (12, 20). OXA-935 (PDB code: 7L5V) belonged to the space group P2_1,_ and OXA-14 (PDB code: 7L5R) belonged to the space group P2_1_2_1_2_1_ (Tables S5, S6). Structural alignment of these revealed a root mean square deviation (r.m.s.d.) of 0.65 driven primarily by a different conformation of the Ω-loop, where the F153S mutation was located (Fig. 3A-C, Fig. S3). In chain A of OXA-14, the Ω-loop was closer to the active site and the indole group from W154 interacted with the carbamylated K70 (3.2 Å) as was seen in OXA-10 (Fig. 3D). The interaction between K70 and W154 is critical for both the activity and the stability of OXA-10 (21). In the chain B of OXA-14, K70 was decarbamylated, causing an open confirmation, consistent with what was observed for OXA-10. In contrast, the Ω-loops of both chains in the crystal structure of OXA-935 had open conformations, the hydrogen bond between K70 and the indole group of W154 was lost (16.4 Å), and both K70 residues were decarbamylated (Fig. 3E, Fig. S3). This more open confirmation of the Ω-loop led to a larger and more positively charged active site cavity, which may allow it to accommodate bulkier and more negatively charged substrates such as ceftazidime (Fig. 3F,G). An additional structure of OXA-935 determined (PDB code: 7N1M) from different crystallization conditions and a different space group, P2_1_2_1_2_1_, also revealed that both monomers of OXA-935 had decarbamylated K70 residues and disordered Ω-loops supporting the observations that S153 conferred significant flexibility to the Ω-loop.

**FIGURE 2.**
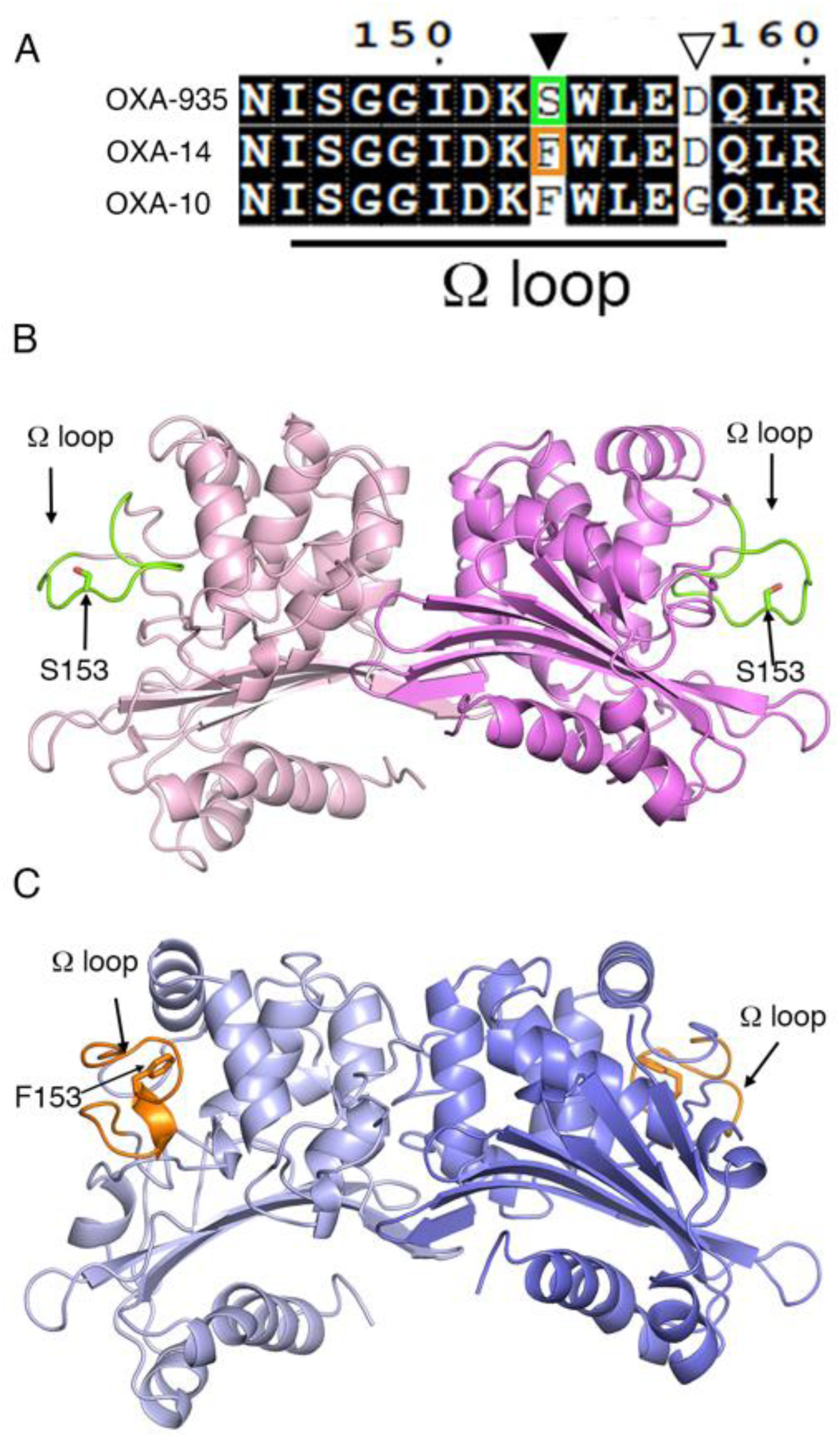
Differences in the structures of OXA-935 and OXA-14. **(A)** Sequence alignment of the Ω-loop of OXA-935, OXA-14 and OXA-10 indicating in orange (F) and green (S) the changes in the residue 153. Cartoon representation of the asymmetric dimeric structures of **(B)** OXA-935 and **(C)** OXA-14 showing the Ω-loop in green for OXA-935 and orange for OXA-14.

**FIGURE 3.**
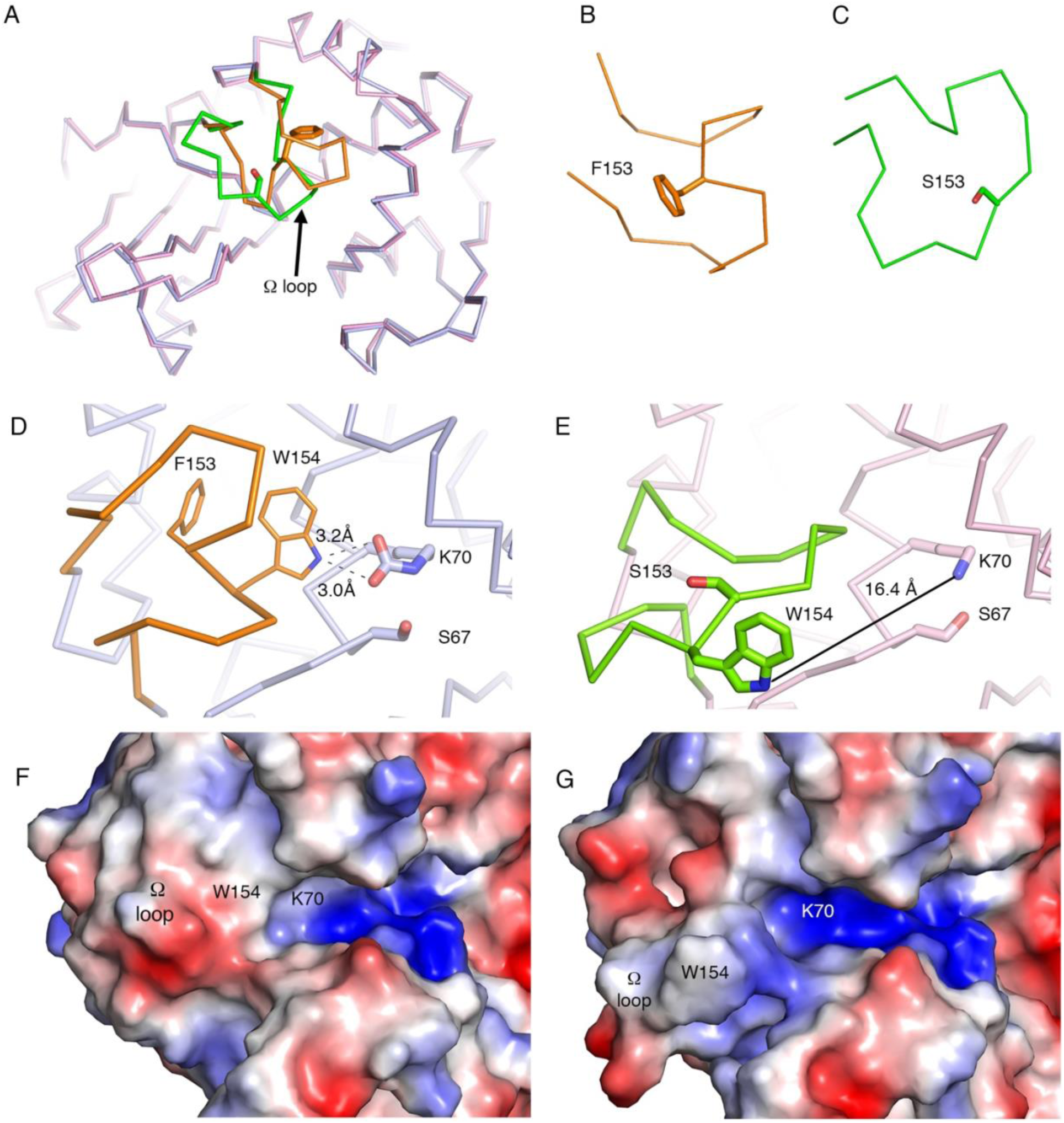
The F153S substitution disrupts the interactions of W154 in the Ω-loop of OXA-935 with the catalytic residue K70. **(A)** Structural alignment of OXA-14 (blue) and OXA-935 (pink) highlighting the Ω-loop in orange and green, respectively. Zoomed in view of the Ω-loop of **(B)** OXA-14 and **(C)** OXA-935. Position of the Ω-loop and interactions of W154 and the catalytic residue K70 in **(D)** OXA-14 and **(E)** OXA-935 Dashed lines represent hydrogen bond interactions. The continuous black line shows the distance between K70 and W154 in OXA-935. Surface charge representation of the Ω-loop and the active site of **(F)** OXA-14 and **(G)** OXA-935.

The carbamylation of K70 in other class D β-lactamases is favored at more basic pH (22, 23), whereas OXA-935 showed decarbamylated K70 even at a pH of 8.3. We hypothesized that the F153S substitution conferred enough flexibility to the Ω-loop to destabilize the carbamylated state of K70 even at basic pH. To corroborate this hypothesis, we used nitrocefin hydrolysis to examine the activity of OXA-14 and OXA-935 across a range of pH values (pH 7.0 −8.5) in a saturated sodium bicarbonate buffer. Sodium bicarbonate stimulated the activity of both enzymes across the tested pH range (Fig. S2). We observed an increase in nitrocefin hydrolysis for OXA-14 as pH values increased suggesting that K70 was more stably carbamylated at higher pH. Conversely, we observed little increase from the baseline in nitrocefin hydrolysis by OXA-935 as pH increased. Thus, unlike OXA-10, OXA-14, and other related β-lactamases, it is likely that K70 of OXA-935 remains decarbamylated *in vitro* despite increases in pH.

### Nitrocefin hydrolysis by OXA-935 is slower than OXA-14

The enzymatic activities of OXA-14 and OXA-935 were tested using nitrocefin as a substrate. Despite the fact that both proteins harbor the G157D mutation and both proteins are responsible for ceftazidime resistance, there were significant differences in their kinetic parameters (Table 3). Consistent with previous reports, the enzyme-substrate binding efficiency of OXA-14 showed cooperativity even at low enzyme concentrations (2.5 nM) and the K_0.5_, the concentration of nitrocefin at which OXA-14 was at half maximum velocity, was 10.1 ± 0.5 µM (7, 20). Contrary to OXA-14, OXA-935 exhibited Michaelis-Menten kinetics and its *K*_*m*_ for nitrocefin (20.8 ± 1.7 µM) was almost double the K_0.5_ of OXA-14, suggesting that the affinity of OXA-935 for nitrocefin was approximately half that of OXA-14. When comparing OXA-935 to OXA-14, we saw an even more dramatic effect on the substrate turnover rate. OXA-935 had a 4.5-fold lower *k*_*cat*_ than the V_max_ of OXA-14 for nitrocefin, suggesting that the F135S mutation present in OXA-935 had a dramatic impact on the ability of OXA-935 to hydrolyze nitrocefin (Table 3). The crystal structure showed that the F135S mutation of OXA-935 led to increased Ω-loop conformational flexibility and a lack of carbamylation at K70 (Fig. 3E) (22). It is conceivable that the lack of carbamylation at the K70 residue led to the poor nitrocefin hydrolysis observed in our biochemical assay. Despite the fact that the nitrocefin hydrolysis kinetics for OXA-935 were slower than for OXA-14, MBD testing confirmed that expression of OXA-935 was sufficient for significant resistance to ceftazidime (Table 2). Additional studies are needed to understand the dynamics of the activation of OXA-935 in the context of our broth dilution assay.

**TABLE 3.** Steady-state kinetic constants of nitrocefin hydrolysis for OXA-14 and OXA-935^a^

|  | *bla*<sub>OXA-14</sub> (G157D) |  | *bla*<sub>OXA-935</sub> (G157D, F153S) |  |
| --- | --- | --- | --- | --- |
|  | *v*<sub>max</sub> (s<sup>-1</sup>) | K<sub>0.5</sub>(μM) | k<sub>cat</sub> (s<sup>-1</sup>) | K<sub>m</sub> (μM) |
| Nitrocefin | 18.5 ± 0.5 | 10.1 ± 0.5 | 4.0 ± 0.169 | 20.8 ± 1.7 |

<sup>a</sup>The data represent the means ± standard deviations from three independent experiments.

## DISCUSSION

In this study, we characterized a novel OXA-10-family β-lactamase, OXA-935, found in three XDR ST-298* *P. aeruginosa* isolates unique with respect to their resistance to ceftazidime. OXA-935 differs from OXA-10 by two amino acid substitutions, G157D which it shares with OXA-14 and F153S which is novel. We showed that deletion of *bla*_OXA-935_ from our three isolates restored susceptibility to ceftazidime and that expression of *bla*_OXA-935_ in laboratory *P. aeruginosa* strains was sufficient for resistance to ceftazidime. We also determined the crystal structures of both OXA-14 and OXA-935. In the crystal structure of OXA-935, the critical active site K70 residue was decarbamylated in both monomers and the F153S substitution resulted in significant flexibility of OXA-935’s Ω-loop. These structural changes likely impact the ability of OXA-935 to hydrolyze bulkier substrates such as ceftazidime.

The most well characterized pathway of antimicrobial resistance to 3^rd^ generation cephalosporins, such as ceftazidime and the newer β-lactam/β-lactamase combination antimicrobials, is the accumulation of chromosomal mutations leading to the overproduction of the intrinsic cephalosporinase AmpC (19). Overexpression of AmpC is often secondary to mutations in the AmpC repressor, AmpD. AmpD inactivation in *P. aeruginosa* results in both moderate basal-level expression and hyperinducible expression of AmpC (14) and is known to lead to extended spectrum beta-lactam resistance. Genome sequencing of our three isolates revealed a mutation in AmpD, resulting in a truncated N-terminus (Δaa 2-30). Despite this AmpD truncation, deletion of *bla*_OXA-935_ restored ceftazidime susceptibility and its ectopic expression produced resistance in laboratory strains of *P. aeruginosa*. This suggests that the resistance phenotype observed in our strains was largely driven by OXA-935 expression and less impacted by the co-existing AmpD truncation. Interestingly, the G157D and F153S substitutions had a more unpredictable impact on the MICs to additional β-lactam compounds other than ceftazidime. When compared to OXA-10, expression of both OXA-14 and OXA-935 resulted in decreased MICs to aztreonam and piperacillin-tazobactam. This suggests that these substitutions, while extending the spectrum of activity against ceftazidime, may negatively impact the ability of OXA-935 to hydrolyze other β-lactam compounds and that there is likely a trade-off to mutations that affect the confirmation of the active site and the flexibility of the Ω-loop.

In *P. aeruginosa* there is growing concern for the increasing spectrum of antimicrobial resistance among OXA-10-family β-lactamases. OXA-10 has been subject to significant mutational pressure both in the laboratory where ceftazidime (6), carbapenem (10, 11, 24), ceftolozane-tazobactam and ceftazidime-avibactam (17, 25, 26) resistant mutants have been selected, and clinically, as evidenced by the sheer diversity of described mutations leading to clinically-significant AMR (7, 16, 27-29). Class D β-lactamases share the same general structure with conserved motifs including an active-site serine 67, a carbamylated lysine (K70), and a stabilizing and invariant tryptophan residue (W154) within the Ω-loop (Fig. S4) (21). In order to better understand the impact of the OXA-935 amino acid substitutions on enzymatic function, we determined the crystal structures of OXA-14 and OXA-935. OXA-14 crystallized as a dimer where only one of the two K70 residues was carbamylated (22, 23). The Ω-loop conformation was maintained and the hydrogen bond between the indole group of W154 and K70 was present. The F153S variant found within OXA-935 yielded a distinctly different structure of the Ω-loop whereby W154 was oriented away from the active site cavity, carbamylation of K70 was not observed, and the active site groove was more positively charged. W154, in addition to stabilizing K70, helps orient K70 so that it has optimal interactions with other residues in the active site (21). Critically, mutations in W154 (W154C), deletion of F153-W154, and duplications of negatively charged residues in this region have been identified in *P. aeruginosa* clinical isolates resistant to ceftazidime-avibactam and ceftolozane-tazobactam (17, 30). In addition, the Ω-loop of OXA-935 was considerably more flexible than in OXA-14. This suggests that both F153 and its neighboring residue W154 may be critical sites of mutation that allow for increased substrate accessibility of the active site.

In assessing functional kinetics, purified OXA-935 hydrolyzed nitrocefin at a slower rate than OXA-14; however, expression during MIC testing was sufficient for ceftazidime resistance. It is possible that decarbamylation of K70 is contributing to this reduced activity against nitrocefin in our biochemical assay. While the lack of carbamylation may offer an explanation for poor nitrocefin hydrolysis, it is unclear what the impact is of this lost post-translational modification on ceftazidime hydrolysis. Several possible explanations exist for why OXA-935 hydrolyzes ceftazidime efficiently despite poor nitrocefin hydrolysis. First, it is possible that OXA-935 may be more highly expressed than OXA-14, more efficiency transported to the periplasm, or more stable once it arrives there. Alternatively, it is possible that carbamylation of K70 is favored under conditions that were disrupted during purification and crystallization. In this case, nitrocefin hydrolysis may be a poor substitute for the kinetics of ceftazidime hydrolysis. Attempts to crystalize OXA-14 or OXA-935 with bound ceftazidime were unsuccessful which may be due to rapid hydrolysis by both enzymes and a lack of K70 carbamylation in OXA-935. Regardless, the new structures of OXA-14 and OXA-935 provide structural insights into the interplay between the Ω-loop and the active site which governs the broad spectrum of activity for this family of enzymes.

Finally, OXA-10-family β-lactamases are often found in mobile transmissible elements such as plasmids (17). OXA-935 is no exception in that it is located within a plasmid-borne integron that harbors additional AMR elements (1). The mobility of OXA-10-family β-lactamases coupled with their propensity for mutation in the face of antibiotic pressure make this enzymatic class one of growing concern. The discovery of yet another variant of OXA-10, OXA-935, that confers ceftazidime resistance, and non-susceptibility to ceftazidime-avibactam and ceftolozane-tazobactam prior to the clinical introduction of the latter two antimicrobial agents is worrisome. Minimal amino acid changes in OXA-10 lead to clinically significant extended-spectrum β-lactamase activity, especially when located in the Ω-loop. Most concerning is the development of resistance to ceftolozane-tazobactam and ceftazidime-avibactam which are often used as last-resort therapy to treat MDR *P. aeruginosa* infections. Ultimately, our findings emphasize the importance of continued molecular surveillance of multidrug-resistant *P. aeruginosa* and increased recognition of the contribution of OXA-10-family β-lactamases to AMR phenotypes.

## MATERIALS AND METHODS

### Bacterial Strains and Growth Conditions

*P. aeruginosa* PS1793, PS1796, and PS1797 are clinical strains from the respiratory tract of patients at Northwestern Memorial Hospital (NMH) isolated between 2005 and 2007 and PABL048 is a clinical strain from the bloodstream of a patient at NMH isolated in 2001 (1). PA14 and PAO1 are commonly used laboratory strains (31, 32). Relevant characteristics of these strains are listed in Table S7.

*Escherichia coli* strain TOP-10 (Invitrogen) was used for cloning and *E. coli* strains S17.1 λpir (33) and SM10 λpir were used to introduce plasmids into *P. aeruginosa. E. coli* BL21(DE3) with and without the pMagic plasmid (34) were used for protein expression. Bacterial strains were streaked from frozen cultures onto LB agar and, unless otherwise stated, grown at 37°C in LB.

Antibiotics were used at the following concentrations: irgasan 5 µg/mL (irg), hygromycin 500 µg /mL (hyg), and gentamicin 100 µg/mL (gent) for *P. aeruginosa*, and gentamicin 15 µg/mL, hygromycin 100 µg/mL, kanamycin 50 µg/mL (kan), and ampicillin 200 µg/mL (amp) for *E. coli*. Further details on the strains and plasmids used in this study can be found in Tables S7 and S8.

### Hybrid assembly of PS1793 and comparison to PABL048

For short-read and long-read sequencing, genomic DNA was extracted from an overnight culture of PS1793 using a Promega Maxwell Cell DNA Purification Kit (Promega Corp., Madison, WI). For short-read sequencing, a sequencing library was prepared using a Nextera XT kit (Illumina, San Diego, CA) and sequenced using an Illumina MiSeq instrument and a v3 flow cell, yielding 2 × 301 bp paired-end reads for a total of 1,174 Mbp of sequence, with an approximate coverage of 160-fold. For long-read sequencing, genomic DNA from PS1793 was used to create a sequencing library using ligation sequencing kit SQK-LSK109 (Oxford Nanopore, UK, catalogue number SQK-LSK109) and sequenced on the Oxford Nanopore MinION platform using a FLO-MIN106 flow cell. Base calling with default quality score filtering and demultiplexing of sequenced reads was performed using Guppy (v3.4.5) yielding 31,050 reads totaling 297 Mbp of sequence and approximate coverage of 41-fold. Hybrid genome assembly was performed using Unicyler (v0.4.8) (35) with default settings to generate a single circular 6,868,713 bp chromosome and 3 circular plasmids totaling 318,215, 113,189, and 69,506 bp respectively. The PS1793 complete genome was annotated using the NCBI Prokaryotic Genome Annotation Pipeline (PGAP) (v5.2) (36, 37) and is available through NCBI BioSample SAMN12162657 and GenBank locus accessions CP083366-CP083369. PS1793 plasmid contigs and pPABL048 were aligned to each other and visualized using BRIG (v0.95) (38).

### Sequencing of PS1796 and PS1797 and comparison to PS1793

For short-read sequencing of PS1796 and PS1797, genomic DNA was extracted from an overnight culture as described previously. Sequencing libraries were prepared using a Nextera XT kit (Illumina, San Diego, CA) and sequenced using an Illumina MiSeq instrument and v3 flow cell as described previously. Reads were quality trimmed and adapter sequences were removed using Trimmomatic (v0.36) (39). Trimmed reads were aligned to the PS1793 complete genome (chromosome and plasmids) using BWA (v0.7.15) (40) (https://arxiv.org/abs/1303.3997v2) and sorted and indexed using samtools (v0.1.19-44428cd) (41). SNV sites were then identified as described previously (1).

### Alignment and phylogenetic comparison of OXA-10-family β-lactamases

For alignment visualization, multiple alignment of all OXA-10 family β-lactamase amino acid sequences was performed using Qiagen CLC Sequence Viewer (v8.0) with default parameters. For phylogenetic analysis, we used all OXA-10 family β-lactamase amino acid sequences and included OXA-5, a relatively closely related class D β-lactamase, as an outgroup protein (42). Sequences were aligned using MUSCLE (v3.8.31) (43). A maximum likelihood phylogenetic tree was constructed based on this alignment using RAxML (v8.2.11) (44) with automatic protein model selection and gamma model of rate heterogeneity (-m PROTGAMMAAUTO) and 1000 rapid bootstraps to assess support (-f a -N 1000). The resulting tree was plotted using iTOL v.6 (45).

### Genus-wide screen for OXA-10-family β-lactamase prevalence

OXA-10 family β-lactamase sequences were obtained from the NCBI Pathogen Detection Reference Gene Catalog using “OXA-10” as the search query in January 2021. OXA-16 was additionally included based on literature review (27). *Pseudomonas* genus genomes available in the *Pseudomonas* Genome Database (46), along with accompanying metadata (species, MLST) were obtained on November 10^th^, 2020. After the exclusion of one genome which failed to download, this yielded a total of 9799 genomes (including both complete and draft genomes). Nucleotide BLAST alignments were performed using each OXA-10 family β-lactamase as the query sequence and each genome as the subject sequence. In order to identify which genomes contained known OXA-10 family β-lactamase genes, BLAST results were parsed to identify alignments with 100% sequence identity and coverage. In order to identify genomes containing any OXA-10 family β-lactamase gene, BLAST results were parsed to identify alignments with 90% sequence identity and coverage for *bla*_OXA-10_.

### Generation of pEX18HygB and *bla*_OXA-14_ and *bla*_OXA-935_ expression plasmids

The hygromycin B resistance gene (*hygR*) and associated promoter (*ampR* promoter) were amplified from the pFLP_hyg (47) plasmid using primers TT115 and TT116, creating a 1,185 bp amplicon. Inverse PCR with primers TT113 and TT114 were used to amplify the backbone of the pEX18Ap plasmid (48), excluding the ampicillin resistance gene and associated promoter, resulting in a 4,644 bp amplicon. Both amplicons were designed with matching overhangs to allow for annealing through Gibson assembly. The *hygR* insert and the inverse PCR product of the pEX18Ap backbone were mixed at a 5:1 ratio and ligated for 30 minutes at 50°C with New England Biolabs (NEB) Gibson Assembly ® Cloning Kit to create pEX18HygB. The resulting assembled product was transformed into chemically competent *E. coli* TOP10 and plated on LB agar plates supplemented with 100 μg/mL hygromycin B (GoldBio, USA). An individual colony was picked and transferred to 5 mL of LB medium supplemented with 100 μg/mL hygromycin B and incubated overnight at 37 °C. After 14 h, pEX18HygB was isolated from *E. coli* TOP10 using a QIAprep Spin Miniprep Kit (Qiagen, Germany) and was verified by sequencing (using 9 pairs of overlapping primers, TT117 – TT125) at the NuSeq facility at Northwestern University. The resulting sequences confirmed the creation of the 5,829 bp plasmid, pEX18HygB.

For the creation of OXA β-lactamase expression vectors for protein purification, both full length (FL) and mature sequence (trunc, Δaa1-20) *bla*_OXA-14_ (NCBI Reference Sequence WP_064056056.1) and *bla*_OXA-935_ (NCBI Reference Sequence WP_141989064.1) were codon optimized (SmartGene®) for expression in *E. coli*, synthesized (Twist Bioscience) and cloned into the pMCSG53 vector (49), which contains a tobacco etch virus (TEV) cleavable N-terminal 6x His-tag, ampicillin resistance and genes for rare codons.

For the creation of plasmids with isopropyl β-d-1-thiogalactopyranoside (IPTG)-inducible expression of *bla*_OXA-10_, *bla*_OXA-14_ and *bla*_OXA-935_, the full length (fl) and mature (tr) sequences without their native promoters were amplified from PABL048 (for *bla*_OXA-10_), from pMCSG53-*oxa14* FL (for *bla*_OXA-14_), and from PS1793 (for *bla*_OXA-935_) using the primers pPSV37_OXA_F_Gibs, pPSV37_OXA_trunc_F_Gibs and pPSV37_OXA_R_Gibs, respectively. HindIII-digested pPSV37 (50) was mixed at a 1:3 ratio with each OXA β-lactamase gene product and ligated using the NEB Gibson Assembly ® Cloning Kit. The resulting vectors, pPSV37-tr-*oxa10*, pPSV37-fl-*oxa10*, pPSV37-tr-*oxa14*, pPSV37-fl-*oxa14* and pPSV37-tr-*oxa935*, pPSV37-fl-*oxa935* were verified by sequencing using primers SeqFwPr and SeqRevPr PSV37 and they, along with pPSV37 vector control, were transformed into electrocompetent (51) PAO1 and PA14. All recombinant methods including introduction of plasmids expressing OXA-10, OXA-14 and OXA-935 in PAO1 and PA14 were reviewed and approved by the Northwestern University Institutional Biosafety Committee. All primer sequences are listed in Supplementary Table S8.

### Generation of Δ*bla*_OXA-935_ *P. aeruginosa* PS1793, PS1796, and PS1797

Upstream and downstream fragments surrounding the *bla*_OXA-935_ gene were amplified from PS1793 genomic DNA using the following primers: oxa10 5-1-HindIII, oxa10 5-2, oxa10 3-1, and oxa10 3-2-HindIII where oxa10 5-2 and oxa10 3-1 contain a 24-bp overlapping linker sequence (*TTCAGCATGCTTGCGGCTCGAGTT*) to generate an in-frame deletion of the *bla*_OXA-935_ gene (Table S8). The resultant upstream and downstream fragments were used as templates for overlap extension PCR and amplified with oxa10 5-1-HindIII and oxa10 3-2-HindIII to create a single linear fragment for insertion. The integration proficient vector, pEX18HygB, was cut with HindIII and the plasmid and insertion fragment were ligated using the NEB Gibson Assembly ® Cloning Kit. The resulting vector, pEX18HygB-Δ*bla*_OXA-935_, was verified by sequencing at the NuSeq facility at Northwestern University and transformed into *E. coli* SM10 λpir. Following conjugation and allelic exchange with PS1793, PS1796 and PS1797 whole-genome sequencing was performed on all mutant strains to confirm the mutation. Briefly, genomic DNA was isolated from an overnight culture of each parental strain and its corresponding mutant, sequencing libraries were prepared using a Nextera XT kit, and sequencing performed using an Illumina MiSeq instrument and a v2 flow cell, yielding 2 × 251 bp paired end reads. Reads were quality trimmed and aligned to the PS1793 complete genome as previously described. The site of the deletion was examined using Tablet v1.21.02.08 (52).

### Protein production and purification

Gene sequences of the mature sequence from OXA-14 and OXA-953 were cloned as described above. The plasmids were transformed into *E. coli* BL21(DE3)(Magic) cells (53) and transformants were cultured in Terrific Broth (TB) medium supplemented with 200 μg/ml ampicillin and 50 μg/ml kanamycin. The expression of the protein was induced by addition of 0.5 mM IPTG when cultures reached OD_600_=1.8-2.0; cultures were further incubated at 25 °C at 200 rpm for 14 h (53). The cells were harvested by centrifugation, resuspended in lysis buffer (50 mM Tris pH 8.3, 0.5 mM NaCl, 10% glycerol, 0.1% IGEPAL CA-630) and frozen at −30°C until purification. Frozen suspensions of bacteria were thawed and sonicated at 50% amplitude, in 5 s x 10 s cycle for 20 min at 4°C. The lysate was centrifugated at 39,000 x *g* for 40 min at 4°C, the supernatant was collected, and the protein was purified as previously described with some modifications (22, 54). The supernatant was loaded onto a His-Trap FF (Ni-NTA) column using a GE Healthcare ÅKTA Pure system in loading buffer (50 mM Na+/Phosphate buffer pH 7.8, 0.5 M NaCl, 1 mM Tris(2-carboxyethyl) phosphine (TCEP) and 5% glycerol). The column was washed with 10 column volumes (cv) of loading buffer followed by 10 cv of wash buffer (50 mM Na+/phosphate buffer pH 7.8, 1 M NaCl, 1 mM Tris(2-carboxyethyl) phosphine (TCEP) and 5% glycerol, 25 mM imidazole), and was eluted with elution buffer (50 mM Na+/phosphate buffer pH 7.8, 0.5 M NaCl, 1 mM Tris(2-carboxyethyl) phosphine (TCEP) and 5% glycerol, 1 M imidazole). The protein was loaded onto a Superdex 200 26/600 column and run with loading buffer. The peak fraction was collected, mixed with TEV protease (1:20 protease:protein) and incubated overnight at room temperature to remove the 6xHis-tag. The cleaved protein was separated from TEV and the tag by affinity chromatography (Ni-NTA) and dialyzed in crystallization buffer (20 mM K^+^/Na^+^ pH 7.8) for 2 h, concentrated to 8-9 mg/ml and set up for crystallization immediately or flash-frozen and stored at −80°C for further use.

### Crystallization and data collection

Purified OXA-14 or OXA-935 proteins were set up as 2 µl crystallization drops (1µl protein:1µl reservoir solution) in 96-well plates (Corning) using commercially available Classics II, Anions and Ammonium sulfate suites (Qiagen). Diffraction quality crystals of OXA-14 apo-form (PDB code 7L5R) grew from 0.1 M bicine pH 9.0, 2.4 M ammonium sulfate and were cryoprotected using 2 M lithium sulfate prior to freezing. Crystals of OXA-935 (PDB code 7L5V) grew from 0.2 M ammonium acetate, 0.1 M Tris pH 8.5, 25% PEG 3350 and crystals for the second OXA-935 structure (PDB code: 7N1M) grew from 0.2 M ammonium iodide, 2.2 M ammonium sulfate (Tables S4, S5) and were similarly cryoprotected.

The data sets were collected at the beam lines 21ID-D and 21ID-F of the Life Sciences-Collaborative Access Team (LS-CAT) at the Advanced Photon Source (APS), Argonne National Laboratory. Images were indexed, integrated and scaled using HKL-3000 (55).

### Structure solution and refinement

The OXA-14 and OXA-935 structures were solved by Molecular Replacement with Phaser (55) from the CCP4 Suite (56) using the crystal structure of the OXA-10 (PDB code 1E3U) as a search model. Initial solutions went through several rounds of refinement in REFMAC v. 5.8.0258 (57) and manual model corrections using Coot (58). The water molecules were automatically generated using ARP/wARP (59) and ligands were manually fit into electron density maps. The Translation–Libration–Screw (TLS) groups were created by the TLSMD server (60) (http://skuldbmsc.washington.edu/~tlsmd/) and TLS corrections were applied during the final stages of refinement. MolProbity (61) (http://molprobity.biochem.duke.edu/) was used for monitoring the quality of the model during refinement and for the final validation of the structure. Final model and diffraction data were deposited to the Protein Data Bank (https://www.rcsb.org/) with the assigned PDB codes: 7L5R (OXA-14), 7L5V (OXA-935) and 7N1M (OXA-935 #2) (Table S6). Structures were visualized using PyMOL v2.4., Schrodinger, LLC®. Composite omit maps were created in CCP4 at the sigma level (62).

### Structural and sequence alignment

The primary amino acid sequences of OXA-10, OXA-14 and OXA-935 were aligned in Clustal Omega (63). The alignment was used to produce the secondary structure depiction using OXA-935 (PDB code 7L5V) as template using ESPript 3.0 server (64). Additional structure alignments were performed using the POSA (65) and FATCAT (66) servers.

### Minimal Inhibitory Concentrations

Minimal inhibitory concentrations (MICs) for *P. aeruginosa* strains were determined in triplicate using the MBD protocol described by Weigand, et al. (67). The following antibiotics were prepared from commercially available sources and were used to assess MICs: piperacillin/tazobactam, cefepime, ceftazidime, ciprofloxacin, meropenem, gentamicin, colistin and aztreonam. For *P. aeruginosa* strains containing expression vectors, 1 mM of IPTG was added to the MIC assay plate to induce OXA-β-lactamase expression.

### Kinetic Assays

The enzyme kinetic parameters of purified OXA-14 and OXA-935 β-lactamases were determined measuring the initial hydrolysis of nitrocefin (λ= 490 nm, and Δε = 17,400 M^-1^cm^-1^) over time using different concentrations of nitrocefin dissolved in 100 mM sodium phosphate with 50 mM sodium bicarbonate, pH 7.0. The reactions were performed using clear bottom 96-well plates (Grenier) at 30°C and measured on the Tecan Safire2 spectrophotometer. The protein concentrations of OXA-14 and OXA-935 were 2.5 nM and 10 nM, respectively. For OXA-935, the *k*_*cat*_ (turnover rate) and *K*_*m*_ (Michaelis constant) values were obtained through nonlinear regression of the data by the Michaelis-Menten equation, using the plot V/[*E*] versus [S] (where V is the initial velocity, and [*E*] and [S] are the enzyme and substrate concentration, respectively). For OXA-14, the initial velocity was plotted against the concentrations of nitrocefin but fitted using the Hill equation to determine K_0.5_ and V_max_. All plots and curve fitting were generated using Prism software (GraphPad v.9.1.2). Each experiment was performed in triplicate.

The effect of carbamylation on OXA-14 and OXA-935 was tested using purified protein (OXA-14 and OXA-935) at 10 nM, incubated with 50 mM of nitrocefin prepared in 100 mM of sodium phosphate buffer at the following pH values: 7.0, 7.5, 8.0, 8.5 with or without 50 mM of sodium bicarbonate as indicated. Solutions were prepared in a 96-well plate with a total volume of 100 µL. Nitrocefin hydrolysis was measured as described above. Each condition was assayed in triplicate. Velocity was determined using the following equation: nitrocefin (µM/min) = ((slope (abs/min)/path length (cm)) x Δε) x 1,000,000. Linear regression of the initial velocity (nitrocefin (µM/min)), was determined and plotted according the pH.

## ACKNOWLEDGMENTS

This work was supported by grants from the National Institutes of Health (NIH)/National Institute of Allergy and Infectious Diseases (NIAID) (R01 AI118257, R01 AI053674, U19 AI135964, K24 104831, and R21 AI129167 awarded to A.R.H), NIH/National Institute of General Medical Sciences (NIGMS) (T32 GM008061 and T32 GM008152 awarded to N.B.P), an American Cancer Society (ACS) Postdoctoral Fellowship (#130602-PF-17-107-01-MPC, awarded to K.E.R.B.) and an ACS Clinician Scientist Development Grant (#134251-CSDG-20-053-01-MPC, awarded to K.E.R.B.). This project has been funded in whole or in part with Federal funds from the NIAID under Contract Nos. HHSN272201299926C and HHSN27201700060C and Grant Nos. U01AI124316 and R01 GM05789 and from the National Institute of Medical Sciences grant GM118187, both from the NIH, Department of Health and Human Services. This work used resources of the Advance Photon Source, a U.S. Department of Energy (DOE) Office of Science User Facility operated for the DOE Office of Science by Argonne National Laboratory under Contract No. DE-AC02-06CH11357. Use of the LS-CAT Sector 21 was supported by the Michigan Economic Development Corporation and the Michigan Technology Tri-Corridor (Grant 085P1000817). This work was also supported by the Northwestern University NUSeq Core Facility and used resources of the Northwestern University Structural Biology Facility which is generously supported by the NCI CCSG P30 CA060553 award to the Robert H. Lurie Comprehensive Cancer Center and was supported in part through the computational resources and staff contributions provided by the Genomics Compute Cluster which is jointly supported by the Feinberg School of Medicine, the Center for Genetic Medicine, and Feinberg’s Department of Biochemistry and Molecular Genetics, the Office of the Provost, the Office for Research, and Northwestern Information Technology. The Genomics Compute Cluster is part of Quest, Northwestern University’s high-performance computing facility, with the purpose to advance research in genomics. The funders had no role in study design, data collection and analysis, decision to publish, or preparation of the manuscript. We would like to thank members of the Center for Structural Genomics of Infectious Diseases (CSGID) and the Hauser, Ozer, and Kociolek laboratories for their valuable comments during numerous discussions of this work.

## Notes

### Competing Interest Statement

The authors have declared no competing interest.

### Summary of Updates

This manuscript version contains supplementary material.

