## Supplementary Figures and Tables for "Functional and Structural Characterization of OXA-935, a Novel OXA-10-family β-lactamase from *Pseudomonas aeruginosa*"

**Supplementary Information**

**SUPPLEMENTARY FIGURES**

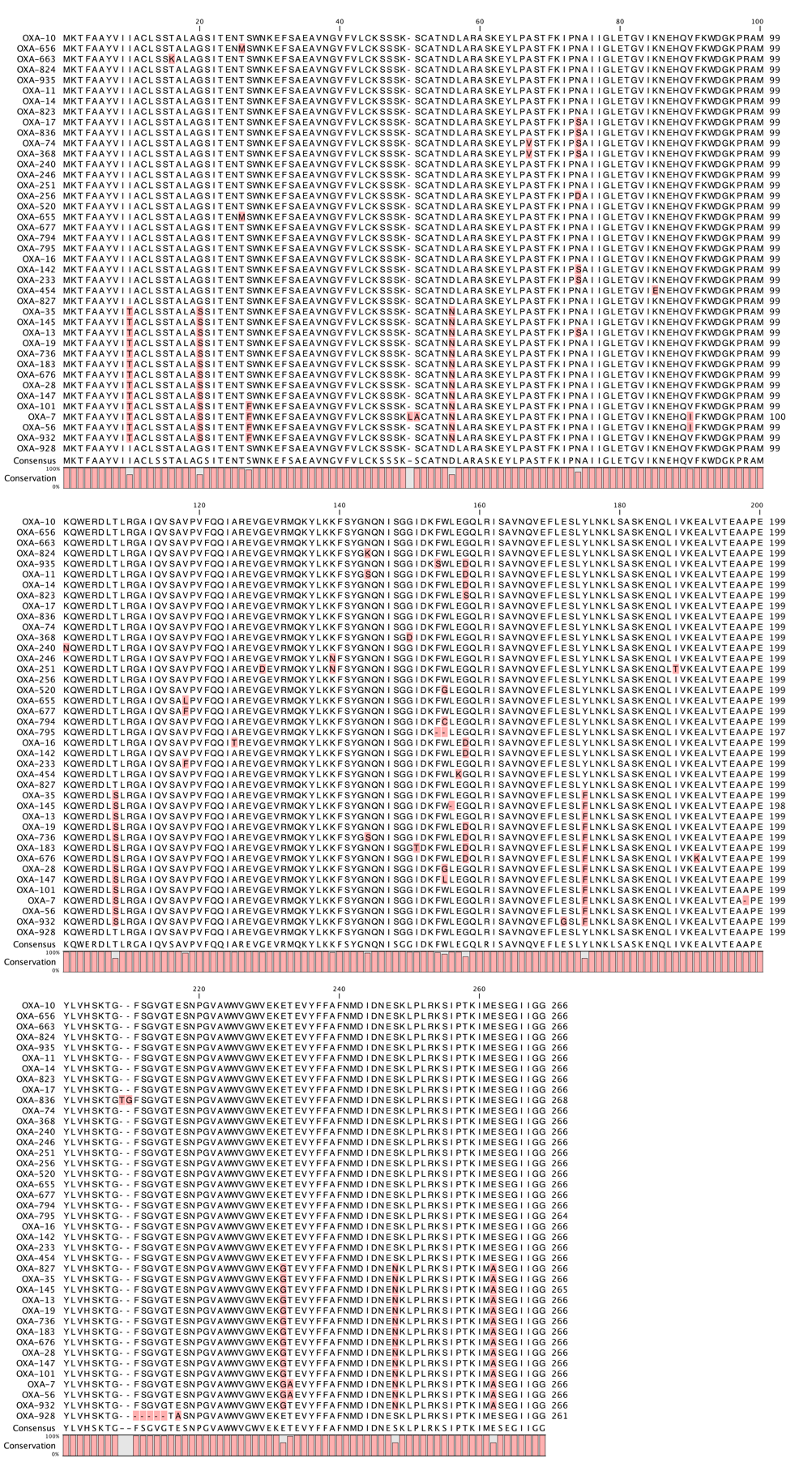

**FIGURE S1.** **Multiple alignment of protein sequences of OXA-10 homologues**. Deviations from the consensus sequence are highlighted in pink.

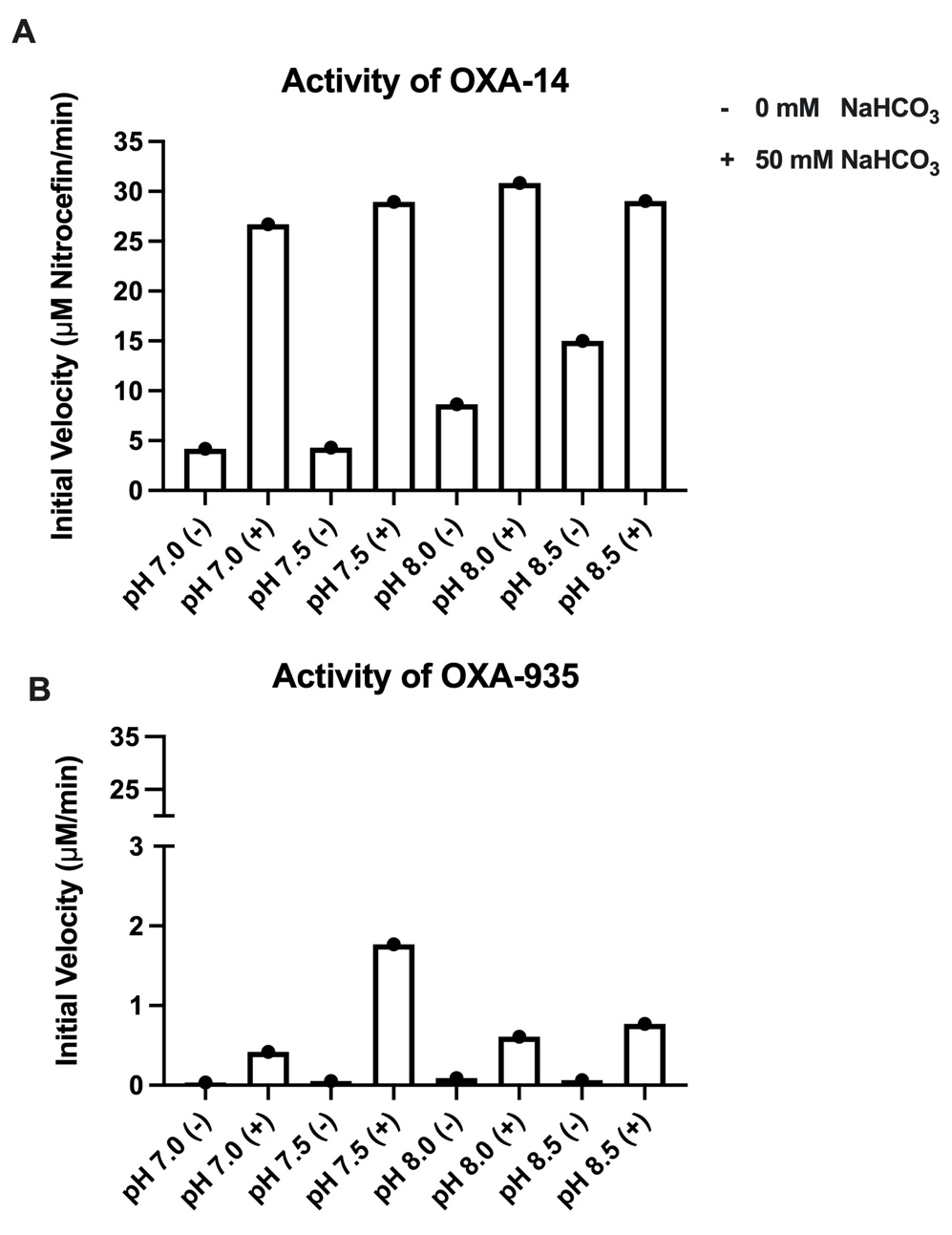

**FIGURE S2. High pH and NaHCO_3_ stimulate the activity of OXA-14.** The plot represents the average initial velocity of nitrocefin hydrolysis of **(A)** OXA-14 and **(B)** OXA-935 in the presence or absence of NaHCO_3_ at different pH values. Experiments were performed in triplicate.

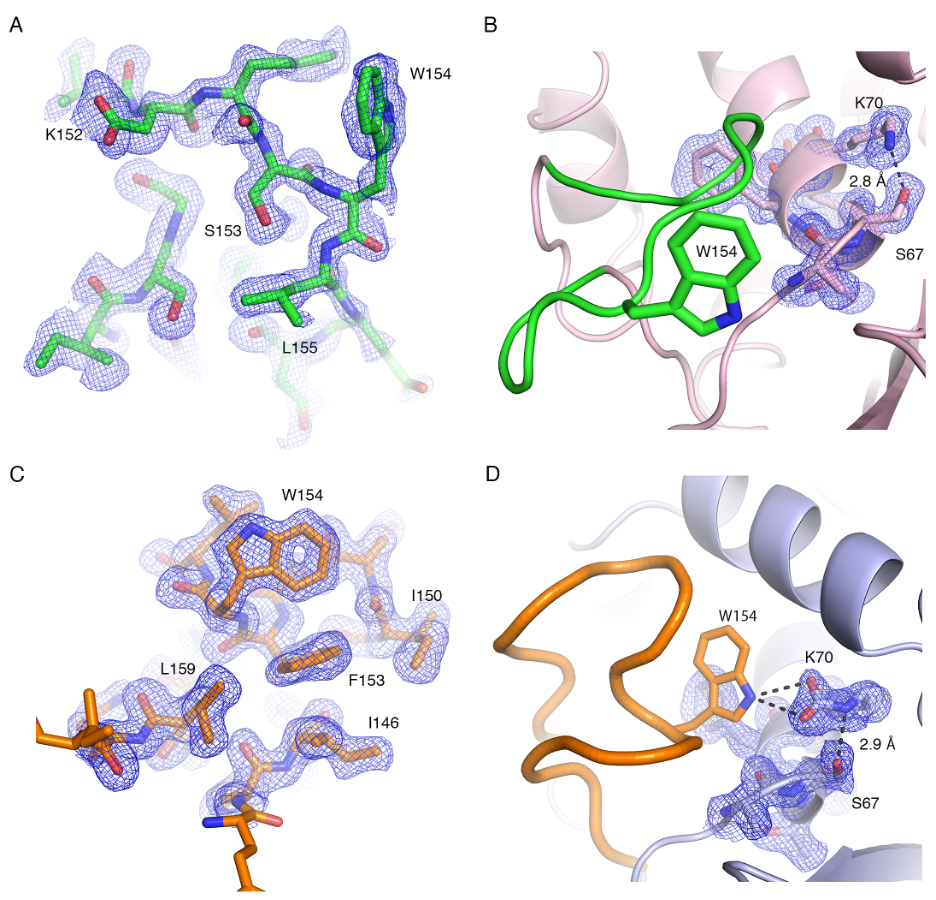

**FIGURE S3. Confirmation of the Ω-loop** **and position of the active site residues in OXA-925 and OXA-14.** Composite omit maps represented as blue mesh surrounding the (**A**) Ω-loop and (**B**) active site residues in OXA-935 and the (**C**) Ω-loop and (**D**) active site residues in OXA-14. Residues are shown as sticks with oxygens in red, nitrogens in blue, carbons in green (Ω-loop, OXA-935), pink (active site, OXA-935), orange (Ω-loop, OXA-14), and pale blue (active site, OXA-14), respectively. Selected residues are labeled and hydrogen bond interactions are shown as dashed lines.

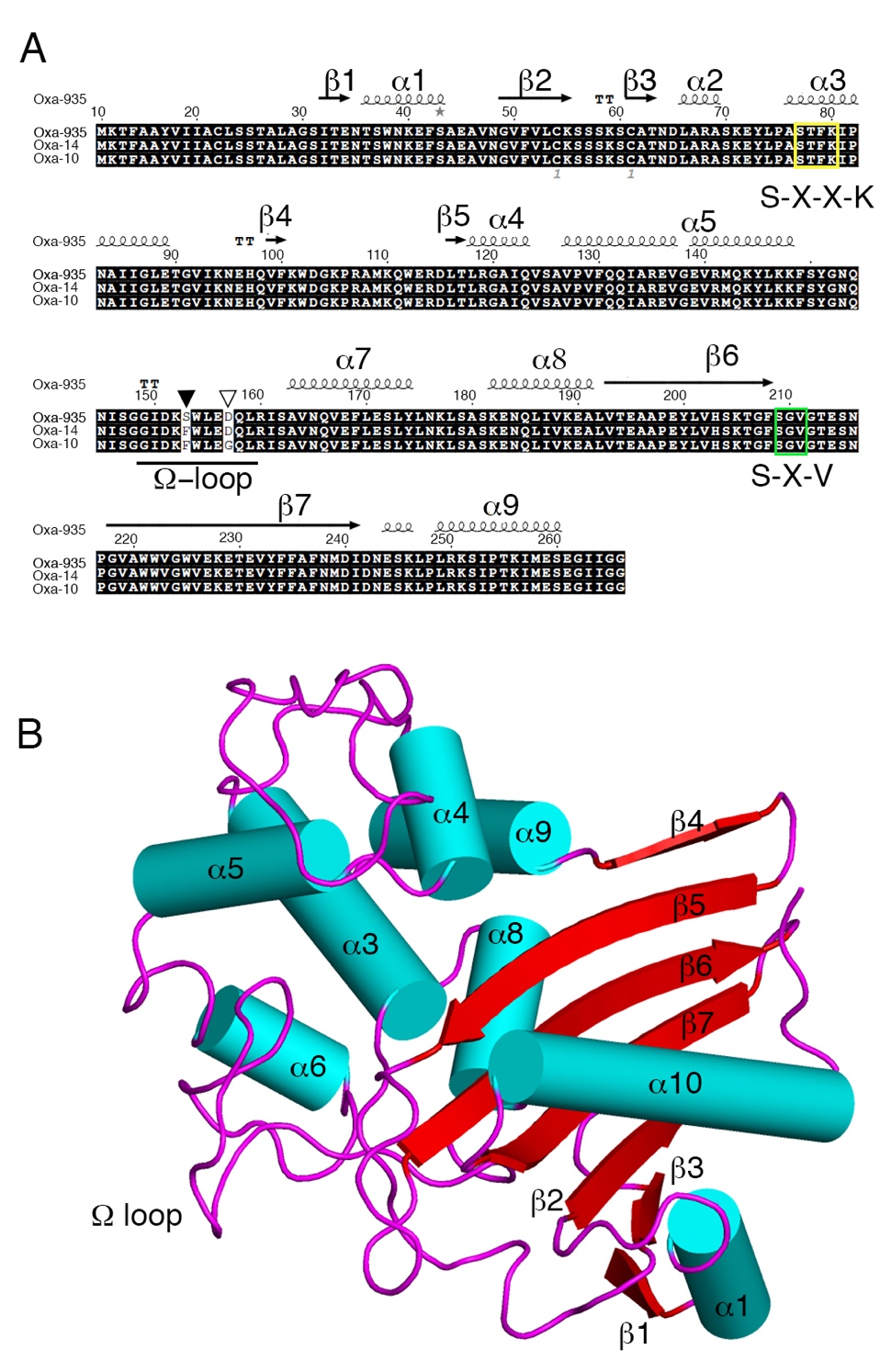

**FIGURE S4. Sequence alignment and structural elements of OXA-10-variants. (A)** Sequence alignment of OXA-935, OXA-14, and OXA-10 where the closed arrowhead highlights the variation at position 153 and the open arrowhead highlights the variation at position 157. Secondary structure elements shown on top of the aligned sequences are based on the OXA-935 structure with α-helices represented as spirals and β-strands as arrows. The catalytic residues, S-X-X-K are highlighted in yellow and S-X-V in green. **(B)** Cartoon representation of the structure of OXA-935, where loops are colored in pink, α-helices are shown as teal cylinders and β-strands as red arrows.

| **TABLE S1** Alignment of PS1796 and PS1797 to the PS1793 Complete Genome | | | | | | | |  |
| --- | --- | --- | --- | --- | --- | --- | --- | --- |
|  | **PS1793 Chromosome** | | **PS1793 p1** | | **PS1793 p2** | | **PS1793 p3** | |
| **Isolate** | **Percent Alignment^a^** | **SNVs** | **Percent Alignment^a^** | **SNVs** | **Percent Alignment^a^** | **SNVs** | **Percent Alignment^a^** | **SNVs** |
| PS1796 | 99.02 | 0 | 99.86 | 0 | 99.84 | 0 | 99.03 | 0 |
| PS1797 | 98.87 | 1 | 99.96 | 0 | 99.65 | 0 | 98.48 | 0 |
| ^a^Percentage of total length of covered by aligned sequences with a minimum depth cutoff of 5 reads. | | | | | | | |  |

| **TABLE S2** Identification of Any OXA-10 Family Gene in *Pseudomonas* Genomes by Species | |
| --- | --- |
| Species | Number of Genomes^ab^ |
| Pseudomonas aeruginosa | 196 |
| Pseudomonas asiatica | 1 |
| Pseudomonas oleovorans | 1 |
| Pseudomonas putida | 2 |
| Pseudomonas sp. | 2 |
| Pseudomonas stutzeri | 8 |
| Pseudomonas xanthomarina | 1 |
| ^a^Based off BLAST search of 9799 *Pseudomonas* genomes with 90% sequence identity and coverage cutoffs | |
| ^b^Only species with at least one OXA-10 family gene identified are displayed | |

| **TABLE S3** OXA-10 Family Genes in *Pseudomonas* by Species and MLST | | |
| --- | --- | --- |
| Gene | Number of Genomes^a^ | Count by Species and MLST |
| *bla*_OXA-10_ | 118 | *Pseudomonas aeruginosa*: 106 (ST233 - 1, ST234 - 7, ST235 - 11, ST244 - 8, ST270 - 1, ST298 - 15, ST308 - 2, ST309 - 5, ST316 - 5, ST357 - 12, ST532 - 1, ST664 - 6, ST708 - 2, ST767 - 1, ST773 - 3, ST1047 - 2, ST1076 - 4, ST1129 - 1, ST1419 - 2, ST2592 - 1, ST2712 - 1, ST3014 - 1, Uknown ST - 14), *Pseudomonas stuzeri*: 8, *Pseudomonas asiatica:* 1, *Pseudomonas olevorans*: 1, *Pseudomonas xanthomarina*: 1, *Pseudomonas sp.*: 1 |
| *bla*_OXA-56_ | 46 | *Pseudomonas aeruginosa*: 46 (ST235 - 1, ST277 - 44, Unknown ST - 1) |
| *bla*_OXA-17_ | 10 | *Pseudomonas aeruginosa*: 10 (ST235 - 7, ST446 - 2, ST639 - 1) |
| *bla*_OXA-101_ | 5 | *Pseudomonas aeruginosa*: 4 (ST111 - 2, Unknown ST - 2), *Pseudomonas asiatica*: 1 |
| *bla*_OXA-19_ | 4 | *Pseudomonas aeruginosa*: 3 (ST235 - 2, ST549 - 1), *Pseudomonas putida*: 1 (Unknown ST - 1) |
| *bla*_OXA-35_ | 4 | *Pseudomonas aeruginosa*: 3 (ST235 - 2, ST348 - 1), *Pseudomonas putida*: 1 (ST70 - 1) |
| *bla*_OXA-935_ | 3 | *Pseudomonas aeruginosa*: 3 (ST298 - 3) |
| *bla*_OXA-74_ | 2 | *Pseudomonas aeruginosa*: 2 (ST235 - 2) |
| *bla*_OXA-14_ | 1 | *Pseudomonas aeruginosa:* 1 (ST235 - 1) |
| *bla*_OXA-28_ | 1 | *Pseudomonas aeruginosa*: 1 (ST235 - 1) |
| *bla*_OXA-246_ | 1 | *Pseudomonas* sp.: 1 |
| ^a^Based off BLAST search of 9799 *Pseudomonas* genomes with 100% sequence identity and coverage cutoffs | | |

| **TABLE S4** Crystallization Information | | | |
| --- | --- | --- | --- |
|  | **OXA-14** | **OXA-935 (monoclinic)** | **OXA-935 (orthorhombic)** |
| PBD code | 75LR | 7L5V | 7N1M |
| Method | Vapor diffusion, sitting drop | Vapor diffusion, sitting drop | Vapor diffusion, sitting drop |
| Plate type | Corning 96-well plate for crystallization | Corning 96-well plate for crystallization | Corning 96-well plate for crystallization |
| Temperature (˚C) | 20 | 20 | 20 |
| Protein Concentration (mg ml^-1^) | 8.0 | 6.0 | 6.0 |
| Protein solution | 0.05 M Sodium phosphate, pH 7.8 | 0.05 M Sodium phosphate, pH 7.8 | 0.05 M Sodium phosphate, pH 7.8 |
| Reservoir solution | 0.1 M Bicine pH 9.0, 2.4 M Ammonium sulfate | 0.2 M Ammonium acetate, 0.1 M Tris pH 8.5, 25% (w/v) PEG 3350 | 0.2 M Ammonium iodide, 2.2 M Ammonium sulfate |
| Volume and ratio of drop | 2 µL (1:1) | 2 µL (1:1) | 2 µL (1:1) |
| Volume of reservoir (μL) | 85 µL | 85 µL | 85 µL |

| **TABLE S5** Data collection and processing | | | |
| --- | --- | --- | --- |
|  | **OXA-14** | **OXA-935** | **OXA-935** |
| Diffraction source | 21ID-F | 21ID-D | 21ID-D |
| Wavelength (Å) | 0.97872 | 1.12713 | 1.12713 |
| Temperature (K) | 100.0 | 100.0 | 100.0 |
| Detector | Rayonix MX-300 | Dectris Eiger 9M | Dectris Eiger 9M |
| Space Group | *P2_1_2_1_2_1_* | *P2_1_* | *P2_1_2_1_2_1_* |
| a,b,c (Å) | 48.81, 96.35, 125.14 | 46.07, 75.04, 82.75 | 47.68, 91.38, 125.79 |
| α, β, γ (˚) | 90.00, 90.00, 90.00 | 90.00, 91.93, 90.00 | 90.00, 90.00, 90.00 |
| Resolution range (Å) | 30.00 – 1.65 (1.68 – 1.65)^a^ | 30.00 – 1.30 (1.32 – 1.30) | 30.00 – 1.95 (1.98 – 1.95) |
| No. of unique reflections | 71,761 (3,530) | 126,515 (5,121) | 40,444 (2,011) |
| Completeness (%) | 100.0 (100.0) | 91.2 (74.0) | 99.8 (100.0) |
| Multiplicity | 6.0 (6.1) | 5.6 (4.9) | 11.1 (10.9) |
| (I/σ(I)) | 27.0 (2.4) | 20.6 (3.2) | 19.4 (2.0) |
| R_p.i.m._ ^a^ | 0.027 (0.365) | 0.033 (0.247) | 0.046 (0.544) |
| CC_1/2_ ^b^ | 0.999 (0.800) | 0.997 (0.854) | 0.994 (0.538) |
| Overall B factor from Wilson Plot (Å^2^) | 21.1 | 10.6 | 33.4 |
| Notes: values in parenthesis are for the outer shell | |  |  |
| ^a^ Estimated Rp.i.m as defined by Weiss, 2001 (1). | |  |  |
| ^b^ Person's correlation coefficient as defined by Karplus and Diedrichs, 2012 (2). | | |  |

| **TABLE S6** Structure refinement | | | |
| --- | --- | --- | --- |
|  | **OXA-14** | **OXA-935** | **OXA-935** |
| Resolution range (Å) | 29.75 – 1.65 (1.69 – 1.65) | 23.95 – 1.30 (1.33 – 1.30) | 29.77 - 1.96 (2.00 - 1.96) |
| Completeness (%) | 99.7 (97.6) | 91.3 (79.4) | 99.4 (95.7) |
| No. of reflections, working set | 68,142 (4,889) | 119,981 (7,667) | 38,401 (2,681) |
| No. of reflections, test set | 3,543 (237) | 6,069 (390) | 1,983 (135) |
| Final R _work_ | 0.164 (0.245) | 0.149 (0.232) | 0.187 (0.296) |
| Final R _free_ | 0.190 (0.259) | 0.170 (0.251) | 0.231 (0.302) |
| No. of non-H atoms |  |  |  |
| Protein | 3,917 | 4,092 | 3,798 |
| Water | 505 | 725 | 170 |
| Ligand | 128 | 0 | 110 |
| R.m.s. deviations |  |  |  |
| Bonds (Å) | 0.006 | 0.005 | 0.006 |
| Angles (˚) | 1.336 | 1.287 | 1.415 |
| Average B factors (Å^2^) | 25.9 | 15.8 | 47.0 |
| Ramachandran plot^c^ |  |  |  |
| Favored regions (%) | 97.0 | 99.0 | 100.0 |
| Additionally allowed (%) | 3.0 | 1.0 | 0.0 |
| Outliers (%) | 0.0 | 0.0 | 0.0 |
| ^c^Validation was done using MolProbity (3). | |  |  |

| **TABLE S7** Bacterial strains used in this study | | | |
| --- | --- | --- | --- |
| **Species** | **Strain ID** | **Relevant Characteristics** | **Reference** |
| *E. coli* | TOP-10 | F- mcrA Δ(mrr-hsdRMS-mcrBC) φ80lacZΔM15 ΔlacX74 nupG recA1 araD139 Δ(ara-leu)7697 galE15 galK16 rpsL(Str^R^) endA1 λ- | Invitrogen |
| *E. coli* | S17-1 λpir | Sm^R^; pro, thi, hsdR^-^M^+^, RP4-2-Tc:Mu;Km:Tn7 λpir | Simon (1983) (4) |
| *E. coli* | SM10 λpir | Km^R^, thi-1 thr leu tonA lacY supE recA::RP4-2-Tc::Mu λpir | Simon (1983) |
| *E. coli* | BL21(DE3)(pMagic) | Km^R^, F- ompT gal dcm lon hsdSB(rB^-^mB^-)^ λ(DE3[lacI lacUV5-T7p07 ind1 sam7 nin5]) [malB^+^]_K-12(_λ^S^), derivative of BL21(DE3) expressing pMagic plasmid which encodes three rare-triplet tRNAs (AGG for Arg, AGA for Arg, and ATA for Ile) | Wu (2000) (5) |
| *P. aeruginosa* | PA14 | human wound isolate | Rahme (1995) (6) |
| *P. aeruginosa* | PAO1 | human wound isolate | Holloway (1955) (7) |
| *P. aeruginosa* | PABL048 | human bacteremia isolate, Assembly: GCA_003411785.2 | Scheetz (2009) (8) |
| *P. aeruginosa* | PS1793 | human respiratory isolate, Assembly: GCA_006704595.1 | Pincus (2020) (9) |
| *P. aeruginosa* | PS1796 | human respiratory isolate, Assembly: GCA_006704575.1 | Pincus (2020) |
| *P. aeruginosa* | PS1797 | human respiratory isolate, Assembly: GCA_006704565.1 | Pincus (2020) |
| *P. aeruginosa* | PS1793∆*oxa935* | Clean deletion of *oxa935* (∆aa5-262) | This study |
| *P. aeruginosa* | PS1796∆*oxa935* | Clean deletion of *oxa935* (∆aa5-262) | This study |
| *P. aeruginosa* | PS1797∆*oxa935* | Clean deletion of *oxa935* (∆aa5-262) | This study |

| **TABLE S8** Plasmids and primers | | |
| --- | --- | --- |
| **Plasmid & Primers** | **Relevant Characteristics or Sequence 5’ – 3’** | **Reference** |
| pFLP-hyg | Hyg^R^; p15a ts ori, expresses FLP recombinase | Huang (2014) (10) |
| pEX18Ap | Amp^R^; allelic exchange vector for making unmarked deletions in *P. aeruginosa* | Hoang (1998) (11) |
| pEX18HygB | Hyg^R^; allelic exchange vector, derivative of pEXT18Ap | This study |
| pEX18HygB-∆*bla_oxa935_* | Hyg^R^; allelic exchange vector for making unmarked deletion of *oxa935* (∆5-262) | This study |
| pMCSG53 | Ap^R^/Cb^R^; ColE1/pBR322/pUC ori+, N-term TEV cleavage site, N-term His_6_ tag | Eschenfeldt (2013)(12) |
| pMCSG53-*oxa14* FL | Ap^R^/Cb^R^; full length version of *oxa14* cloned in frame into the SspI site | This study |
| pMCSG53-*oxa935* FL | Ap^R^/Cb^R^; full length version of *oxa935* cloned in frame into the SspI site | This study |
| pMCSG53-*oxa14* trunc | Ap^R^/Cb^R^; *oxa14* ∆aa1-20 (signal sequence) cloned in frame into the SspI site | This study |
| pMCSG53-*oxa935* trunc | Ap^R^/Cb^R^; *oxa935* ∆aa1-20 (signal sequence) cloned in frame into the SspI site | This study |
| pPSV37 | Gm^R^, ColE1/pMB1/pBR322/pUC ori+, *lacI^q^*, and the *lac*UV5 promoter, derivative of pPSV35 | Lee (2010) |
| pPSV37-*oxa10* | Gm^R^, full length *oxa10* cloned into HindIII site in MCS of pPSV37 | This study |
| pPSV37-*oxa14* | Gm^R^, full length *oxa14* cloned into HindIII site in MCS of pPSV37 | This study |
| pPSV37-*oxa935* | Gm^R^, full length *oxa935* cloned into HindIII site in MCS of pPSV37 | This study |
| TT113 | ATTAGCTTCAAAAGCGCTCTCGTGAGTTTTCGTTCCACTGA | This study |
| TT114 | TGCAGCGAATTGGGGATCTTGAAAACCTCTGACACATGCAG | This study |
| TT115 | AGAGCGCTTTTGAAGCTAATCCCGACGCACTTTGCAGATCT | This study |
| TT116 | AAGATCCCCAATTCGCTGCAGAGTTTTCGCCCCGAAGAACG | This study |
| TT117 | TGCAGCGAATTGGGGATCTTG | This study |
| TT118 | AGATTCTTCGCCTTGGTAGCC | This study |
| TT119 | CTTGTTTTGCAAACTTTTTGA | This study |
| TT120 | GGTTTCATCAGCCATCCGCTT | This study |
| TT121 | ACTCTCGCATGGGGAGACCCC | This study |
| TT122 | AGCAAAAGGCCAGCAAAAGGC | This study |
| TT123 | CCGCTCATGAGACAATAACCC | This study |
| TT124 | ACTTCGAGCGGAGGCATCCGG | This study |
| TT125 | CCCCAGCACTCGTCCGAGGGC | This study |
| oxa10 5 up | TGGCGTTGCGTATGCTCACAG | This study |
| oxa10 3 down | CGAGTCCCGACACCAGACTGCAT | This study |
| oxa10 5-1-HindIII | GTAAAACGACGGCCAGTGCCAGTCTCTCGCGCTCAAGT | This study |
| oxa10 5-2 | *AACTCGAGCCGCAAGCATGCTGAA*AAATGTTTTCATGGCACCTTG | This study |
| oxa10 3-1 | *TTCAGCATGCTTGCGGCTCGAGTT*ATTGGTGGCTAAAACAAAGTT | This study |
| oxa10 3-2-HindIII | GTCGACCTGCAGGCATGCAGGCTTCATGCTTCACG | This study |
| pPSV37_OXA_F_Gibs | CGACCTGCAGGCATGCAATGAAAACATTTGCCGCATATG | This study |
| pPSV37_OXA_R_Gibs | CCAAGGGGTTATGCTAATTAGCCACCAATGATGCCC | This study |
| pPSV37_OXA_trunc_F_Gibs | CGACCTGCAGGCATGCAATGTCAATTACAGAAAATACGTCTTG | This study |
| SeqFwPr PSV37 | GCGATCAAAAAACCCCTCAA | This study |
| SeqRevPr PSV37 | CAGGAAACAGCTATGACCAT | This study |
